# Rbbp8nl contributes to resistance to anticancer immunotherapy by creating a non-inflamed tumor microenvironment and attenuating CD8+ T cell infiltration

**DOI:** 10.1101/2024.01.21.575810

**Authors:** Zhenglin Yi, Jinhui Liu, Hao Deng, Jiao Hu, Xiongbing Zu

## Abstract

Immune checkpoint blockade (ICB) monotherapy has limited efficacy, and it is crucial to explore predictive markers for the efficacy of immunotherapy and its use as new therapeutic targets or sensitization targets in combination with immunotherapy. Pan-cancer research using RNA sequencing data from The Cancer Genome Atlas (TCGA) revealed that RBBP8NL is particularly overexpressed in the tumor microenvironment (TME) of different malignancies. Additionally, there is a negative correlation between RBBP8NL and immune modulators, T cell inflammatory scores, immunological checkpoints, cancer immune cycle, and tumor-infiltrating immune cells (TIICs), so it can be inferred that RBBP8NL shapes the non-inflammatory TME in BLCA. BLCA patients with high RBBP8NL expression have a low response rate to immunotherapy, and they are more likely to experience hyperprogression of the illness. Interestingly, anti-RBBP8NL with immunotherapy could work better together than they do separately. According to the results of the multi-omics analysis, RBBP8NL inhibits the recruitment of cytotoxic lymphocytes. Furthermore, RBBP8NL’s predictive significance for immunotherapy success has been confirmed across several immunotherapy cohorts. In summary, RBBP8NL is a crucial component of the TME and a developing target for binding to ICB as well as a biomarker to direct precision medicine.

## 1. Introduction

Bladder cancer (BC) is one of the most common malignancies of the genitourinary system. Globally, there are approximately 430,000 newly diagnosed BC cases and more than 165,000 BC patient deaths every year (1). BC can be classified as either muscle-invasive bladder cancer (MIBC) or non-muscle-invasive bladder cancer (NMIBC) based on the depth of tumor invasion (MIBC). NMIBCs make up around 70% of newly diagnosed bladder cancers. Nevertheless, even when the bladder tumor is removed transurethrally, 30% of patients experience a recurrence and develop invasive bladder cancer (2).

Lately, through the development and use of immune checkpoint inhibitors (ICI), the safety and effectiveness of immunotherapy in advanced bladder cancer have been confirmed (3). ICI has been suggested by guidelines for the treatment of metastatic or locally advanced bladder cancer that is incurable (4). Although immune checkpoint blockade (ICB) provides survival benefits to some patients with advanced bladder cancer. However, a lot of individuals continue to have acquired or primary resistance, making them nonresponsive to ICB monotherapy (5). For ICB to be effective, an inflammatory TME and preexisting anticancer immunity are required. Theoretically, molecules or pathways that cause the TME to be non-inflamed would lead to resistance to ICBs (6). Therefore, it is crucial to identify a key target that can transform the non-inflammatory TME into an inflammatory TME and explore its potential as a combined sensitization target for immunotherapy.

RBBP8NL (RBBP8 N-Terminal Like) is a Protein Coding gene. Located in extracellular space. Silencing RBBP8NL in normal human primary keratinocytes (NHPK) resulted in a significant increase in HSV-1 replication. RBBP8NL-silenced NHPK reduced IFNk gene expression but increased IL1b. In addition, the keratinocyte differentiation markers keratin 10 (KRT10) and loricrin (LOR) were similarly shown to have reduced gene expression when RBBP8NL was silenced (7). Research has revealed that the RBBP8NL gene is related to the progression and prognosis of MIBC and is a central gene in the development of bladder cancer (8). Nevertheless, the function of RBBP8NL in regulating the TME status of bladder cancer has not yet been explored.

We conducted a pan-cancer examination of RBBP8NL’s expression patterns and immunological consequences in this work, and we screened out RBBP8NL which might result in non-inflammatory TME in BLCA. We also carried out single-cell RNA sequencing to investigate potential molecular processes and found that RBBP8NL functions by inhibiting the recruitment of cytotoxic T lymphocytes (CTLs). Our research indicates that BLCA might be a good fit for anti-RBBP8NL treatment. In addition, we also find that RBBP8NL has the capacity to forecast BLCA molecular subtypes and immunotherapy efficacy.

## 2. Methods

### Data extraction and preprocessing

The UCSC Xena data portal was used to get the Cancer Genome Atlas (TCGA) data (9). To compute the tumor mutational burden (TMB), log2 converts the RNA-seq data and evaluates the somatic mutation data using VarScan2. Microsatellite instability (MSI) data were gathered from Bonneville’s research (10). To obtain copy number variation (CNV) data, utilize the UCSC Xena data portal, and the LinkedOmics data site to acquire methylation data (11). The expression data for RBBP8NL in normal tissues were obtained from the Genotypic Tissue Expression (GTEx) and the BioGPS data Portal. Lastly, consult the Cancer Cell Line Encyclopedia (CCLE) to learn more about RBBP8NL expression in cancer cell lines.

### Assessment of the TME’s immunological properties in BLCA

The TME in BLCA showcases immunological traits such as immunomodulator expression, cancer immunity cycle activity, tumor-infiltrating immune cells (TIICs) infiltration levels, as well as immunological checkpoint expression inhibition. Our initial data collection focused on 122 immunomodulators, encompassing MHC, receptors, chemokines, and immune stimulators, as reported in Charoentong et al.’s study (12). The anticancer immune response is reflected in the cancer immunity cycle, which involves seven sequential steps: the release of cancer cell antigens (Step 1), presentation of cancer antigens (Step 2), priming and activation (Step 3), immune cell trafficking to tumors (Step 4), immune cell infiltration into tumors (Step 5), recognition of cancer cells by T cells (Step 6), and killing of cancer cells (Step 7) (13). The fate of tumor cells is determined by the activities of these steps. Xu and colleagues evaluated their endeavors by means of single-sample gene set enrichment analysis (ssGSEA), which is predicated on the gene expression levels of distinct samples (14).

Subsequently, various algorithms were created to assess the infiltration levels of TIICs in the TME through the analysis of bulk RNA-seq data. Calculation mistakes may arise from the use of distinct marker gene sets and methods for TIICs. To mitigate these errors, we conducted a comprehensive assessment of TIIC infiltration levels using 7 distinct algorithms: Cibersort-ABS, MCP-counter, quanTIseq, TIMER, xCell, TIP, and TISIDB (15–20). To verify the impact of RBBP8NL on cancer immunity modulation in BLCA, an analysis of the relationship between RBBP8NL and the TME’s immunological features was done, specifically addressing the aspects mentioned above. The results obtained from this investigation underwent validation in 3 distinct external cohorts.

### RNA sequencing of specimens with BLCA

60 fresh samples from BLCA patients were procured at Xiangya Hospital and promptly preserved in liquid nitrogen. The extraction of total RNA from the tissues was carried out using TRIzol (Invitrogen, Carlsbad, CA, USA) under the manufacturer’s guidelines. The Agilent 2100 bioanalyzer and NanoDrop were then used to quantify total RNA (Thermo Fisher Scientific, MA, USA). The construction of the mRNA library ensued, involving the purification and fragmentation of whole RNA into tiny fragments. The synthesis of first-strand cDNA and second-strand cDNA was subsequently carried out, after the incubation with an A-tailing mix and RNA Adapter Index for end repair, PCR amplification of the cDNA fragments was performed.

The final library, which is represented by single-stranded circular DNA, was constructed using the qualifying double-stranded PCR products that were obtained. Out of the sixty initially gathered samples, 57 met the qualification criteria. Subsequently, the 57 qualified bladder cancer samples, designated as the Xiangya cohort, underwent sequencing on the BGISEQ-500 platform (BGI-Shenzhen, China). RSEM was used to calculate the levels of gene expression (v1.2.12).

### Real-time quantitative PCR (qPCR)

Using an RNA extraction reagent (Donghuan, Shanghai, China) in accordance with the manufacturer’s recommendations, total RNA was extracted from thirty matched samples of bladder cancer and nearby normal tissues for the real-time qPCR analysis. An instrument called a cDNA synthesis kit was used to synthesize cDNA (Takara, Dalian, China). Subsequently, real-time qPCR was conducted to assess RBBP8NL’s expression, employing SYBR Green qPCR Master Mix (Junxing, Suzhou, China). Normalization of gene expression levels utilized the GAPDH housekeeping gene. The following primer sequences were used for RBBP8NL and GAPDH: RBBP8NL (forward primer: 5′-GAGATCCACGAGAAGGAAGTCC-3′; reverse primer: 5′-CTTCAGTGTCTTCTGCTGTTCC-3′); GAPDH (forward primer: 5′-CAAGGCTGTGGGCAAGGTCATC-3′; reverse primer: 5′-GTGTCGCTGTTGAAGTCAGAGGAG-3′).

### Enrichment score calculations for different gene signatures

We compiled a collection of gene signatures that exhibit a positive correlation with the clinical reaction to an anti-PD-L1 agent in BLCA (21). Additionally, 12 distinct BLCA signatures associated with distinct molecular subtypes were gathered from the research conducted by the Bladder Cancer Molecular Taxonomy Group (22). In addition to that, various therapeutic signatures were gathered, encompassing oncogenic pathways capable of influencing a non-inflamed tumor microenvironment (TME), gene signatures associated with targeted therapies, and those predicting responses to radiotherapy. The enrichment scores for these signatures were determined using the GSVA R package (23). Following this, the observation was made that the mutation statuses of many pivotal genes, served as factors that indicate how well BLCA patients may respond to neoadjuvant chemotherapy (24–27).

Upon contrasting the variations in enrichment score values for therapeutic signatures and neoadjuvant chemotherapy predictions’ mutation statuses among RBBP8NL groups, we were able to discern the predictive role of RBBP8NL as a reaction to these treatments. To identify genes linked to BLCA as therapeutic targets, the Drugbank database was used for our screening (28).

### Forecasting the molecular subtypes in BLCA

Various molecular subtypes have been established, including CIT, Lund, MDA, TCGA, Baylor, UNC, and Consensus subtypes (29–34). The determination of individual molecular subtypes was carried out utilizing the R packages BLCAsubtyping and ConsensusMIBC. Subsequently, we examined the correlation between RBBP8NL and distinct molecular subtypes, as well as specific gene profiles for BLCA. Building upon the previously illustrated relationships between several molecular subtypes, bladder cancer can be categorized into 2 principal subtypes: Subtypes that are basal and luminal (22). We generated receiver operating characteristic (ROC) curves to assess the forecast precision of RBBP8NL for molecular subtypes. Furthermore, the validation of RBBP8NL’s predictive accuracy was conducted in 4 cohorts for external validation. These cohorts encompassed 2 general BLCA cohorts (GSE69795, GSE48276), 1 cohort in connection with immunotherapy (IMvigor210), and another associated with neoadjuvant chemotherapy (GSE70691).

### Single-cell RNA sequencing

Single-cell suspensions of immune cells were acquired by dissociating subcutaneous tumors from three mice per group and loaded onto a 10X Genomics to produce nanoliter-scale gel emulsion beads (GEMs). Using the Chromium Single Cell 3’ v3 Reagent Kit, barcoded single-cell RNA sequencing (scRNA-seq) libraries were prepared. The GEM facilitated the creation of barcoded full-length cDNA through a reverse transcription reaction. Subsequently, the barcoded full-length cDNA underwent building of libraries, involving processes such as fragmentation, end repair, A-tailing, ligation with tag adapters, and PCR amplification. The MobiNova-100 platform (MobiDrop, Zhejiang, China) was employed for the construction and sequencing of scRNA-seq libraries, following the instructions provided by OE Biotech Go., Ltd. (Shanghai, China).

### Processing of unprocessed data and quality assurance

Cells displaying less than one thousand UMIs or more than twenty percent transcripts from mitochondria were deemed inadequate quality and excluded. Every analysis that followed was carried out using the R environment (version 4.1.2) using Seurat (version 4.3.0). Given that libraries were separately constructed for each sample, sample IDs were employed to mitigate potential batch effects in Harmony (35). Following principal component analysis (PCA), the first 30 principal components (PCs) were utilized for UMAP analysis. Clusters identified through Seurat’s FindClusters function were labeled based on recognized markers provided in the outcomes.

### Analytical statistics

Correlations among variables were examined using either Pearson or Spearman coefficients. With continuous variables between binary categories that follow a normal distribution, a t-test was employed; otherwise, the test of Mann-Whitney U was utilized. The Fisher’s exact test or the chi-squared test were used to compare categorical variables. Utilizing the Kaplan-Meier technique, prognostic assessments of categorical factors were performed to make survival curves, with statistical significance assessed through the log-rank test. For every test, a significance criterion of P < 0.05 was established, and these were two-sided. Version 4.1.2 of the R program was used to conduct the statistical analysis.

## 3. Results

### Analyzing the pan-cancer expression pattern, prognostic relevance, and immunological associations of RBBP8NL

Following an extensive examination of the expression data from the Oncomine, GTEx, and TCGA databases, our findings indicated elevated expression of RBBP8NL in numerous cancers, including bladder cancer and breast cancer, in comparison to normal tissues (Figure S1A-B). Additionally, RBBP8NL demonstrated expression in diverse cancer cell lines, including those related to bladder cancer, as observed through the examination of expression data from CCLE databases (Figure S1C). In a set of 30 matched samples comprising both bladder cancer and normal tissues, the expression of RBBP8NL was notably higher in cancerous tissues compared to normal ones (Figure S1D). The observed pan-cancer overexpression pattern of RBBP8NL prompted an investigation into its prognostic significance. Consequently, we carried out an extensive investigation of pan-cancer survival, taking into account cancer-specific survival, progression-free survival, and overall survival. As expected, RBBP8NL become a predictive biomarker for a number of different malignancies (Figure S2-S4). Nevertheless, it is crucial to note that these results require further validation, particularly through multivariable analysis.

Analyzing the immunological role of RBBP8NL across multiple cancers is crucial for identifying potential candidates for anti-RBBP8NL immunotherapy. Our investigation unveiled a negative correlation between RBBP8NL and numerous immunomodulators in BLCA (Figure 1A). To assess the TME’s immune cell infiltration, we employed the ssGSEA algorithm, revealing a negative relationship between RBBP8NL and several tumor-infiltrating immune cells in BLCA (Figure 1B). Additionally, our findings indicated a mutual exclusivity between the expression of RBBP8NL and numerous immunological checkpoints in BLCA, including as PD-L1, PD-1, CTLA-4, and LAG-3 (Figure 1C-F). Beyond BLCA, the observed adverse immunological relationships between RBBP8NL were not evident in other cancers. However, it was observed that RBBP8NL displayed a negative correlation with tumor mutation burden (TMB) and microsatellite instability (MSI) in several cancers, hinting at a potential role for RBBP8NL as a marker reflecting cancer immunogenicity in these instances (Figure S5).

**Figure 1.**
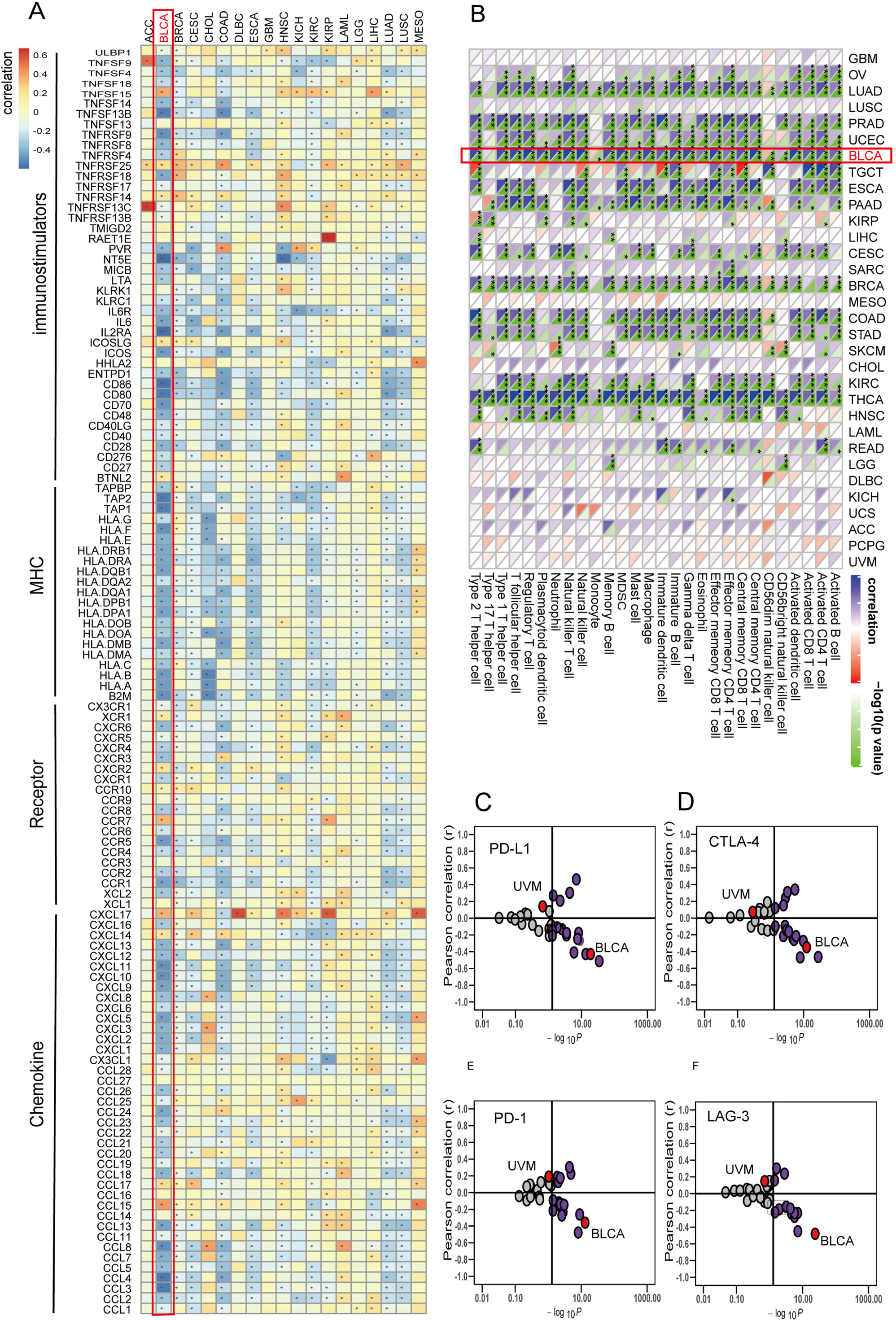
The impact of RBBP8NL on the pan-cancer immune state. (A) Examination of the correlation between RBBP8NL and 122 immunomodulators, encompassing chemokines, receptors, MHC, and immunostimulators. (B) Evaluation of the correlation between RBBP8NL and 28 tumor-associated immune cells computed via the ssGSEA algorithm. The correlation coefficient is color-coded, with asterisks denoting statistically significant p-values derived from Spearman correlation analysis (*P < 0.05; **P < 0.01; ***P < 0.001). (C-F) Analysis of the correlation between RBBP8NL and four immune checkpoints—PD-L1, CTLA-4, PD-1, and LAG-3. Cancer types are represented by dots, with the Y-axis indicating the Pearson correlation and the X-axis reflecting -log10P.

In conclusion, the expression pattern of RBBP8NL appears to be specific to the tumor microenvironment (TME), emphasizing the prospect of RBBP8NL as a focal point for personalized cancer immunotherapy. The pronounced immunosuppressive impact of RBBP8NL in the TME is particularly evident in bladder cancer, suggesting the potential candidacy of bladder cancer for anti-RBBP8NL immunotherapy.

### Analysis of RBBP8NL mutations in BLCA

No mutations were identified in the RBBP8NL gene. The copy number variation (CNV) pattern of RBBP8NL is illustrated in Figure S6A. Significantly, both copy number deletion and methylation of RBBP8NL were associated with reduced expression of RBBP8NL mRNA (Figure S6B-C). These findings suggest that epigenetic modifications of the RBBP8NL gene could serve as an alternative therapeutic intervention for anti-RBBP8NL inhibitors. Figures S6D-F succinctly summarize the top thirty mutated genes in the low and high RBBP8NL groups, as well as the mutation profile of BLCA.

### RBBP8NL forms a non-inflammatory TME in BLCA

A substantial negative correlation was observed between RBBP8NL and numerous immunomodulators (Figure 2A). In the high-RBBP8NL group, a majority of MHC molecules exhibited downregulation, suggesting a decreased capacity for antigen presentation and processing. Additionally, three pivotal chemokines (CXCL9, CXCL10, and CCR3), known to facilitate the recruitment of CD8+ T cells into the tumor microenvironment (TME) in bladder cancer, were decreased in the group with elevated RBBP8NL. Several other chemokines exhibited a negative correlation with RBBP8NL. These chemokines and receptors play a role in promoting the recruitment of effector TIICs. Given the intricate and varied functions of the chemokine system, relying solely on the association of RBBP8NL with individual chemokines falls short of fully understanding the comprehensive immunological influence of RBBP8NL in the tumor microenvironment (TME).

**Figure 2.**
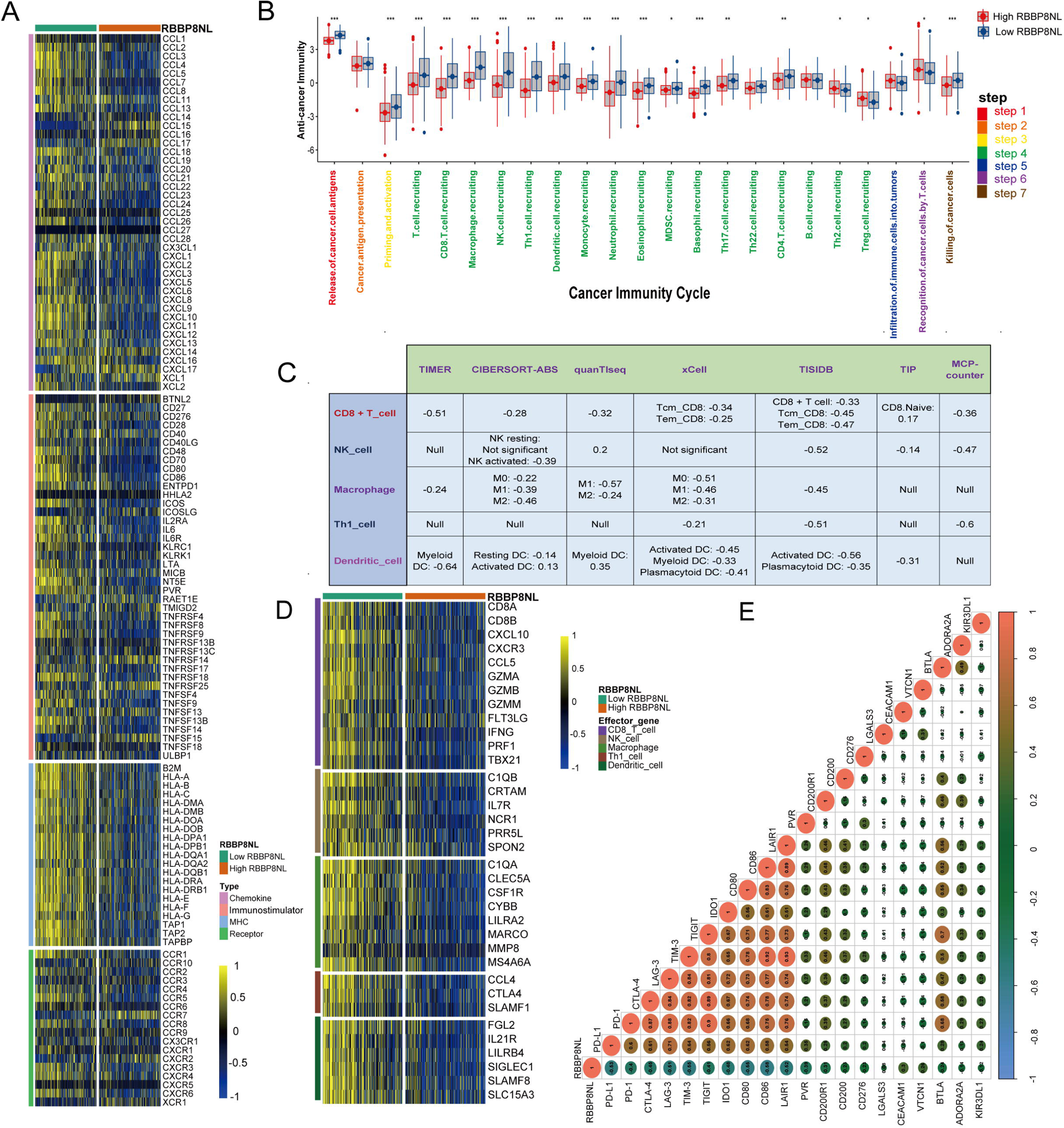
In BLCA, RBBP8NL forms a non-inflammatory TME. (A) Assessment of expression disparities in 122 immunomodulators (chemokines, receptors, MHC, and immunostimulators) between the high- and low-RBBP8NL groups in BLCA. (B) Evaluation of distinctions in the various steps of the cancer immunity cycle between the high- and low-RBBP8NL groups. (C) Investigation into the correlation between RBBP8NL and the infiltration levels of five types of tumor-infiltrating immune cells (CD8+ T cells, NK cells, macrophages, Th1 cells, and dendritic cells), was computed through seven independent algorithms. (D) Identification of differences in the effector genes of the aforementioned tumor-associated immune cells between the high- and low-RBBP8NL groups. (E) Analysis of the correlation between RBBP8NL and 20 inhibitory immune checkpoints. The Spearman correlation coefficient is represented by color and values, with asterisks denoting statistically significant p-values determined using the Mann-Whitney U test (*P < 0.05; **P < 0.01; ***P < 0.001).

The cancer immunity cycle’s functional representation mirrors the combined functionality of both the chemokine system and diverse immune modulators (13, 14). In the high-RBBP8NL group, a reduction in activities related to most steps in the cycle was noted. Encompassing the liberation of cancer cell antigens (Step 1), initiation and activation (Step 3), and the migration of immune cells to tumors (Step 4), notably CD8+ T cell recruitment, Macrophage recruitment, Th1 cell recruitment, NK cell recruitment, DC recruitment, and TH17 recruitment (Figure 2B). Consequently, the diminished activities of these steps have the potential to lower the infiltration levels of effector TIICs in the TME. Notably, the activity related to the recognition of cancer cells by T cells (Step 6) was found to be downregulated in the low-RBBP8NL group, possibly attributed to the elevated expression of PD-L1 in this group. Moreover, it was noted that the high-RBBP8NL group’s actions related to Step 7 (eliminating cancer cells) were constrained.

We then determined the infiltration level of tumor-infiltrating immune cells (TIICs) using 7 distinct algorithms (Figure S7-S13). Consistent with earlier findings, RBBP8NL displayed a negative correlation with the infiltration levels of CD8+ T cells, NK cells, Th1 cells, macrophages, and dendritic cells across diverse algorithms (Figure 2C). Additionally, RBBP8NL exhibited a negative correlation with the effector genes associated with these TIICs (Figure 2D). Significantly, molecules linked to immune checkpoint inhibitors exhibit reduced expression in non-inflammatory tumor microenvironments (36). In our investigation, a consistent negative correlation was observed between RBBP8NL and a majority of immune checkpoint inhibitors (Figure 2E). Overall, this strong correlation suggests a link between RBBP8NL and the promotion of a non-inflamed tumor microenvironment (TME).

### Clinical outcome and ICB hyperprogression in BLCA are predicted by RBBP8NL

Theoretically, heightened expression of RBBP8NL in patients is expected to result in a diminished response to immune checkpoint blockade (ICB), given that RBBP8NL characterizes a non-inflamed tumor microenvironment (TME). Additionally, a negative correlation was observed between RBBP8NL and the pan-cancer T-cell inflamed score (R = −0.53, P < 0.01) (Figure 3A-B). The potential incidence of ICB-related hyperprogression was increased in the high RBBP8NL group. Notably, the high-RBBP8NL group exhibited significantly elevated rates of copy number amplification and mRNA expression for genes positively associated with hyperprogression (Figure S14A-B). Conversely, in the high-RBBP8NL group, there was a significant decrease in the mRNA expression of genes adversely linked to hyperprogression. (Figure S14A-B).

**Figure 3.**
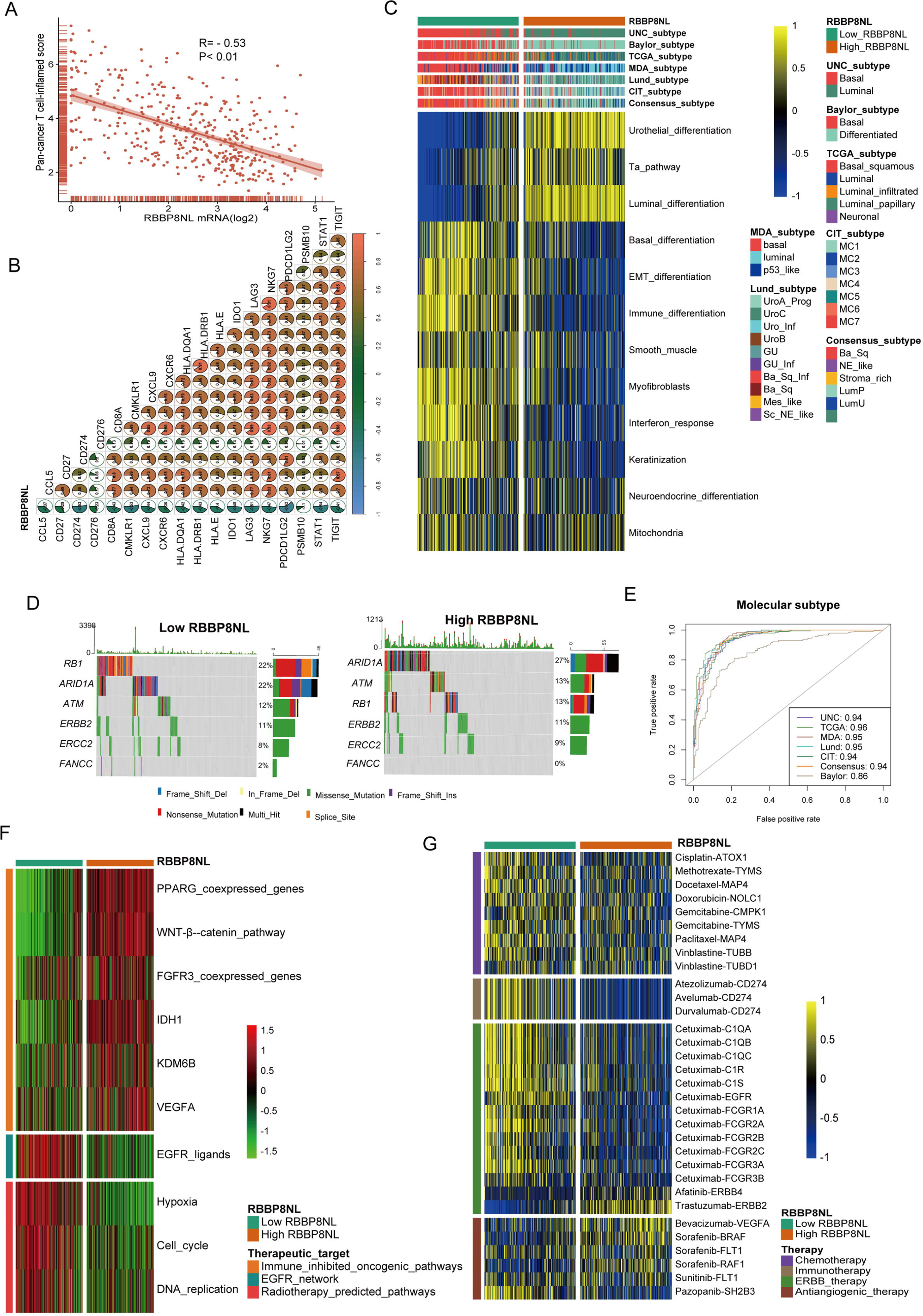
In BLCA, RBBP8NL forecasts the therapeutic response to many medications as well as the molecular subtype. (A-B) Associations between RBBP8NL and the pan-cancer T cell inflamed score and the individual genes comprising the T cell inflamed signature. The T cell inflamed score exhibits a positive correlation with the clinical response to cancer immunotherapy. (C) Relationships between RBBP8NL and molecular subtypes were assessed using seven distinct algorithms and bladder cancer signatures. (D-E) Mutational profiles of neoadjuvant chemotherapy-related genes in the low- and high-RBBP8NL groups. (E) Predictive precision of RBBP8NL for molecular subtypes using seven different algorithms, with accuracy determined by the area under the ROC curves. (F) Correlations between RBBP8NL and the enrichment scores of various therapeutic signatures, including targeted therapy and radiotherapy. (G) Correlation between RBBP8NL and BLCA-related drug-target genes identified from the Drugbank database.

To sum up, implementing immune checkpoint blockade in bladder cancer patients with elevated RBBP8NL expression may not be advisable, as they show reduced responsiveness to ICB and an increased likelihood of hyperprogression.

### Molecular subtypes and potential therapeutic targets are predicted by RBBP8NL

Results from the PURE-01 research provided insights that basal-type bladder cancer displayed the most substantial infiltration of immune cells and higher rates of pathological reaction to ICB drugs (37). Likewise, the subtype analysis arrived at an analogous outcome, emphasizing that basal-type tumors exhibited a greater likelihood of responding to immune checkpoint blockade (ICB) (22). Notably, BLCA with lower RBBP8NL expression tended to align with the basal subtype among the 7 molecular subtypes (Figure 3C). This reaffirmed the observation of a negative correlation between RBBP8NL and the response to ICB. Additionally, higher enrichment scores for luminal differentiation, Ta pathway, and urothelial differentiation were observed in the high-RBBP8NL group. In contrast, the high-RBBP8NL group exhibited lower enrichment scores for basal differentiation, EMT differentiation, immune infiltration, and interferon response (Figure 3C). Furthermore, aside from the Baylor molecular subtyping schemes, RBBP8NL demonstrated an area under the curve (AUC) of ≥ 0.90 within different schemes (Figure 3E). Identical outcomes were noted through validation utilizing 4 cohorts (Figure S14C-F).

A molecular subtype is a valuable predictor of the medical reaction to various treatment modalities (22, 38). Notably, basal subtype malignancies demonstrated a higher likelihood of responding to neoadjuvant chemotherapy. The low-RBBP8NL group, associated with the basal subtype, exhibited significantly elevated mutation rates in RB1, ERBB2, and FANCC (Figure 3D). Furthermore, the low-RBBP8NL group exhibited increased enrichment scores for pathways predicted for radiotherapy and EGFR ligands. (Figure 3F). Additionally, information from the Drugbank database suggests that individuals with low RBBP8NL levels exhibit markedly improved responses to chemotherapy, immunotherapy, and ERBB treatment. (Figure 3G). These findings suggest that immune checkpoint blockade, chemotherapy, and ERBB treatment could be considered, either individually or in combination, for the therapy of bladder cancer with low RBBP8NL expression.

Bladder cancer with elevated RBBP8NL expression tended to align with the luminal subtype (Figure 3C). Notably, immune checkpoint blockade, chemotherapy, and radiotherapy were deemed inappropriate for bladder cancer characterized by high RBBP8NL expression. The high-RBBP8NL group exhibited significantly increased enrichment scores for a number of oncogenic pathways that reduce immunity (Figure 3F). These pathways were intricately associated with the non-inflamed tumor microenvironment in bladder cancer. Therefore, blocking these pathways emerged as a strategy to foster the development of an inflamed TME, consequently resuming anti-cancer immunity (39, 40). In line with this finding, medications that specifically target the FGFR and PPARG pathways have shown encouraging results in the treatment of bladder cancer. For instance, erdafitinib, an FGFR inhibitor, exhibited great responses in advanced bladder cancer following prior immune checkpoint blockade treatment (41). In theory, RBBP8NL and these carcinogenic pathways have similar immunosuppressive roles. Hence, a targeted treatment that inhibits these pathways could be employed in conjunction with anti-RBBP8NL treatment for treating bladder cancer with elevated RBBP8NL expression. Additionally, our findings suggest that anti-angiogenic treatment might be a viable option for bladder cancer characterized by high RBBP8NL expression (Figure 3G).

### Molecular subtypes and immunological characteristics in the Xiangya cohort are predicted by RBBP8NL

Activities related to most steps in the cancer immunity cycle were found to be downregulated in the high-RBBP8NL group (Figure 4A). Within our cohort, RBBP8NL exhibited a negative correlation with most of immunomodulators. Moreover, RBBP8NL discovered to be inversely connected to CD8+ T cells, NK cells, dendritic cells, and macrophages (Figure 4B). As anticipated, RBBP8NL displayed negative correlations with crucial steps of the cancer-immunity cycle (Figure 4C). Additionally, we examined the associations between RBBP8NL and forecasted immune checkpoint blockade response signatures. RBBP8NL exhibited negative correlations with the ratings of enrichment for every favorable signature associated with immunotherapy (Figure 4D). Additionally, RBBP8NL was also inversely connected to the T cell-inflamed score (Figure 4E).

**Figure 4.**
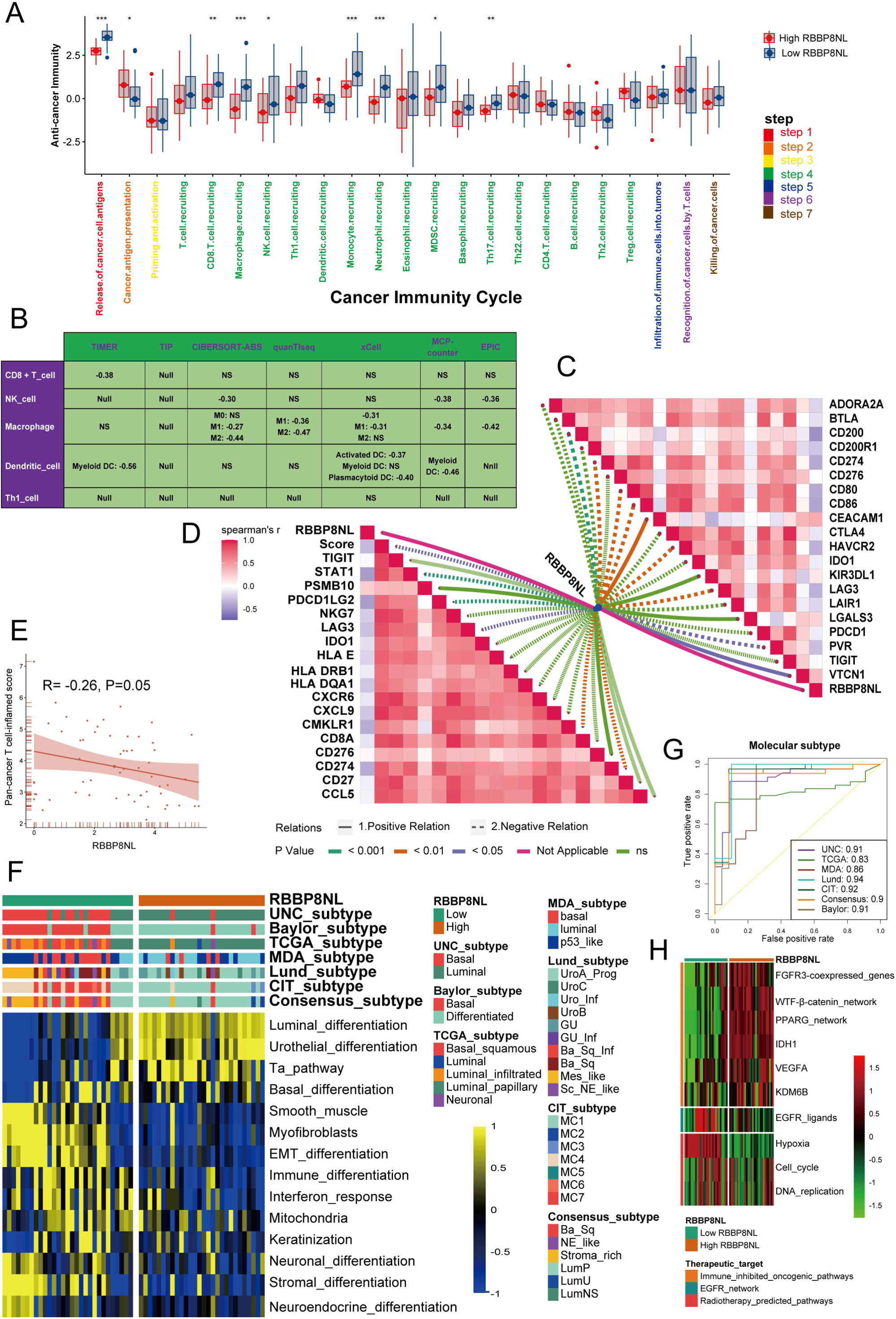
Functions of RBBP8NL in the Xiangya cohort immunological phenotype prediction. (A) Discrepancies in the steps of the cancer immunity cycle between high- and low-RBBP8NL groups. (B) Associations between RBBP8NL and the infiltration levels of five tumor-associated immune cells (CD8+ T cells, NK cells, macrophages, Th1 cells, and dendritic cells). (C) Relationships between RBBP8NL and the steps of the cancer immunity cycle. (D) Links between RBBP8NL and the enrichment scores of immunotherapy-predicted pathways. (E) Correlations between RBBP8NL and the T cell inflamed score. (F) Associations between RBBP8NL and molecular subtypes, as well as bladder cancer signatures. (G) ROC curves illustrate the predictive accuracy of RBBP8NL in anticipating molecular subtypes. (H) Correlations between RBBP8NL and the enrichment scores of therapeutic signatures, encompassing radiotherapy and targeted therapy.

In brief, RBBP8NL exhibits precise differentiation between basal and luminal subtypes in 7 diverse molecular subtype algorithms, as depicted in Figure 4F-G. Bladder cancer characterized by elevated RBBP8NL expression tends to align with the luminal subtype. The predictive accuracy of RBBP8NL for molecular subtypes ranges from 0.83 to 0.94 across the 7 algorithms. Moreover, the validated contributions of RBBP8NL in anticipating the degree of therapeutic response to radiation, targeted treatment, and chemotherapy are effectively demonstrated in Figure 4H.

### Tumor microenvironment modifications are mediated by RBBP8NL inhibition, according to single-cell sequencing

Nevertheless, the constraints inherent to bulk RNA sequencing lie in its characteristics. Gene expression levels are determined by averaging across all cells within the tissue. To address this limitation, we conducted scRNA-seq analysis on 3 bladder cancer samples, examining over 19,000 cells based on well-established biomarkers. The classification of cells into 6 categories—epithelial cells, fibroblast cells, T/NK cells, endothelial cells, B cells, and myeloid cells—was achieved through this approach (Figure 5A). It is essential to highlight that the predominant expression of RBBP8NL is observed in epithelial cells as opposed to immune and endothelial cells. By relying on the accumulated copy number variations (CNV), EPCAM+ epithelial cells are identified as urothelial cells with malignancy. To confirm the expression pattern of RBBP8NL, we incorporated 2 external scRNA cohorts into the analysis. In the GSE135337 scRNA cohort, where over 36,000 cells were analyzed and categorized into 5 clusters, the primary cell subtype expressing RBBP8NL remained malignant bladder urothelial cells (Figure 5B). Therefore, it can be inferred from the majority of RBBP8NL expression patterns on malignant cells that RBBP8NL plays an immunomodulatory role in the TME mainly by changing the characteristics of cancer cells.

**Figure 5.**
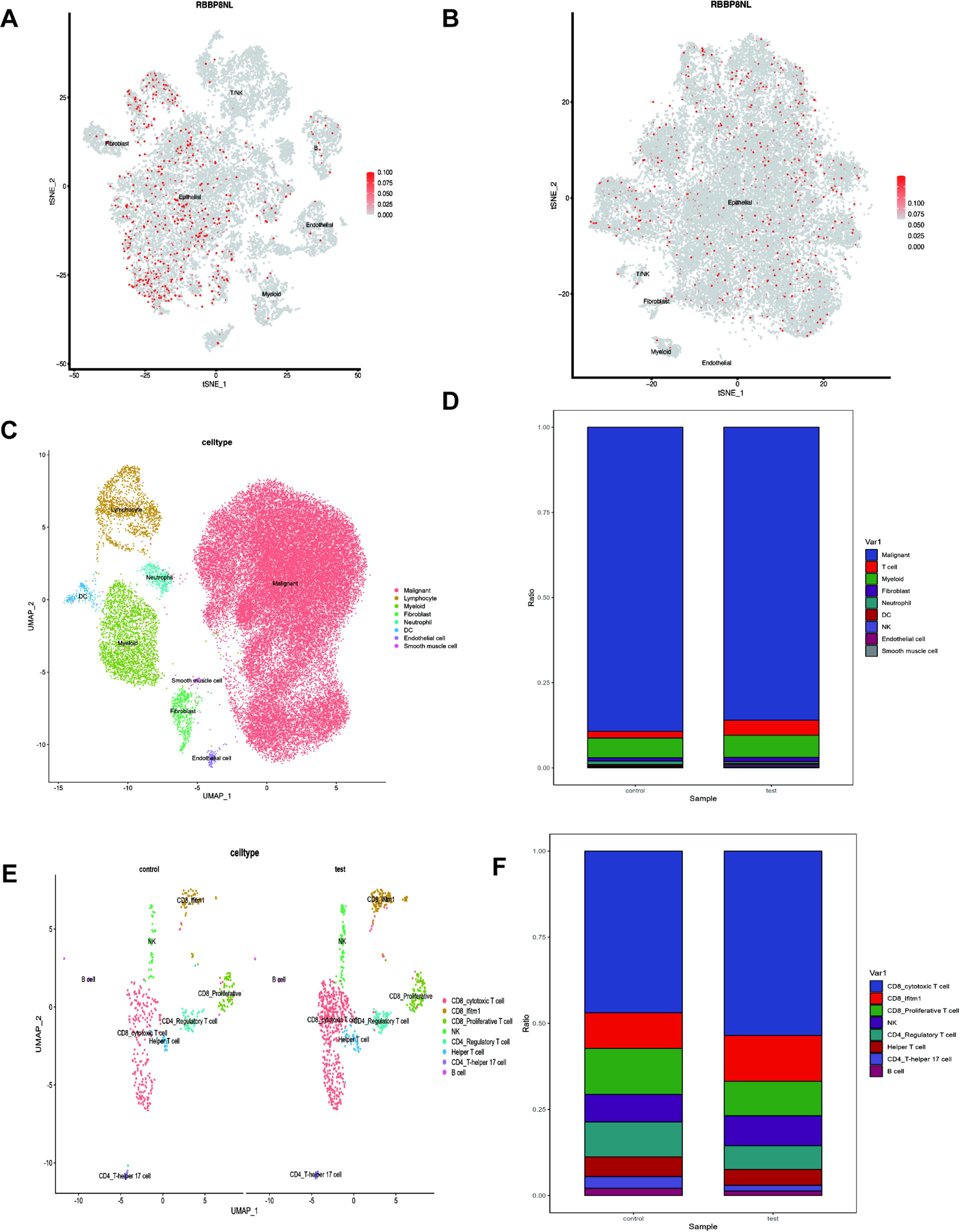
RBBP8NL’s role in forming TME is being investigated using scRNA-seq. (A-B) tSNE plots depicting the single-cell expression pattern of RBBP8NL in the Xiangya scRNA cohort and GSE135337 cohort. (C) Clustering of single cells from subcutaneous tumors (3 tumors from 3 RBBP8NL KD mice vs 3 tumors from 3 control mice) of the control and RBBP8NL KD groups into 8 major cell types. (D) No discernible difference in the proportion of cell clusters between the control and RBBP8NL KD groups. (E) Clustering of lymphocytes from the control group and the RBBP8NL KD group into 8 clusters based on classical markers. (F) Proportions of different lymphocyte subpopulations in the control group and the RBBP8NL KD group (3 tumors from 3 RBBP8NL KD mice vs 3 tumors from 3 control mice). Significance was considered at P-values < 0.05: *P < 0.05; **P < 0.01; ***P < 0.001.

To investigate the effect of RBBP8NL on the in vivo tumor immune microenvironment, we generated a successful RBBP8NL knockout (RBBP8NL KD) mouse (MB49) bladder cancer cell line and established a subcutaneous bladder cancer model using both RBBP8NL KD and control mouse cell lines. At the designated time, tumors were harvested for subsequent analysis. Seeking a comprehensive understanding of the overall alterations in tumor-infiltrating immune cells following RBBP8NL KD, we conducted single-cell RNA sequencing on tissues derived from RBBP8NL KD and control mice. Following quality control and utilizing cell type-specific gene markers, we identified 42,452 high-quality cells that were grouped into 8 main cell types, including lymphocytes, myeloid cells, neutrophils, dendritic cells (DCs), endothelial cells, fibroblast cells, smooth muscle cells, and malignant cells (Figure 5C). However, no substantial changes in the main cellular components were detected between the RBBP8NL KD group and the control group. There was just a little rise in the percentage of T cells (Figure 5D). These findings suggest that RBBP8NL KD might enhance the profusion of various immune cell types, potentially transforming “cold” tumors into a more “hot” tumor microenvironment.

To delve deeper into the T cell alterations influenced by RBBP8NL KD, lymphocytes were categorized into 8 clusters, encompassing CD8-cytotoxic T cells, CD8-Ifitm1, CD8-Proliferative T cells, NK cells, CD4-Regulatory T cells, Helper T cells, CD4-T-helper 17 cells, and B cells (Figure 5E). As depicted in Figure 5F, the most significant change induced by RBBP8NL KD was the upsurge in circulating CD8+ T cells, specifically CD8-cytotoxic T cells. Additionally, a slight reduction in Treg abundance and a modest rise in the percentage of NK cells were observed. Collectively, these findings suggest that RBBP8NL KD activates cytotoxic T cells (CTL) and fosters their expansion, enhancing the cytotoxicity of circulating CD8+ cells and thereby exerting anti-tumor effects.

## 4. Conclusions

This investigation indicates that bladder cancer holds promise as a viable candidate for anti-RBBP8NL immunotherapy. Our findings demonstrate that RBBP8NL contributes to a non-inflammatory tumor microenvironment in bladder cancer and can effectively forecast the clinical response to immune checkpoint blockade and bladder cancer molecular subtypes. Leveraging single-cell sequencing, we present a thorough overview of the alterations in the tumor microenvironment orchestrated by RBBP8NL inhibition. Inhibiting RBBP8NL elicits anti-tumor effects by augmenting the percentage of tumor-infiltrating NK cells and circulating CD8+ T cells. Simultaneously, our study unveils a synergistic sensitization strategy involving RBBP8NL inhibition and ICB, offering novel biomarkers to guide the precision treatment of BLCA.

## Supporting information

Supplementary figures and their legends.

## Declarations

### Ethics approval and consent to participate

Approval for this study was granted by the Ethics Committee at Xiangya Hospital, Central South University. Each participant provided informed consent by signing the necessary documentation.

### Availability of data and materials

The datasets utilized or analyzed in the present study can be obtained from the corresponding author upon reasonable request.

### Competing interests

The authors declare that they have no competing interests.

### Funding

This work was supported by the National Natural Science Foundation of China (81873626, 81902592), Hunan Natural Science Foundation (2020JJ5884), Hunan Province Key R&D Program (2019SK2202) and Xiangya Hospital Youth Fund (2018Q09).

### Authors’ contributions

HJ and ZXB conceived and supervised the study. YZL and DH contributed to data collection and assembly. HJ, YZL, and LJH performed data analysis and interpretation. YZL, LJH, and DH performed software, visualization, and validation. All authors contributed to writing the manuscript. All authors reviewed and approved the final manuscript.

## Acknowledgments

We express our sincere gratitude to all individuals who participated in this study.

## Notes

### Competing Interest Statement

The authors have declared no competing interest.

