## Supplementary figures and their legends. for "Rbbp8nl contributes to resistance to anticancer immunotherapy by creating a non-inflamed tumor microenvironment and attenuating CD8+ T cell infiltration"

**Supplementary files**

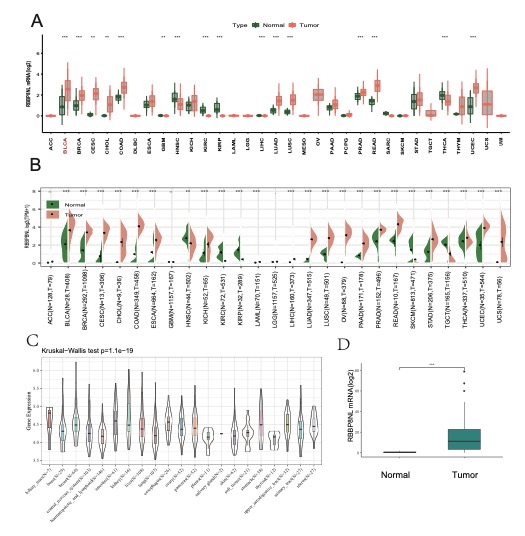

**Figure S1. The immunological connection and expression pattern of RBBP8NL among pan-cancers.** (A-B) Expression patterns of RBBP8NL across pan-cancers in TCGA, TCGA combined with GTEx, and Oncomine. Significant statistical p-values were denoted with asterisks and calculated using the Mann-Whitney U test. (*P < 0.05; **P < 0.01; ***P < 0.001). (C) RBBP8NL expression in cancer cell lines from CCLE. (D) Quantitative RT-PCR (qPCR) assessment of Siglec-15 mRNA levels in 30 paired bladder cancer and normal tissues.

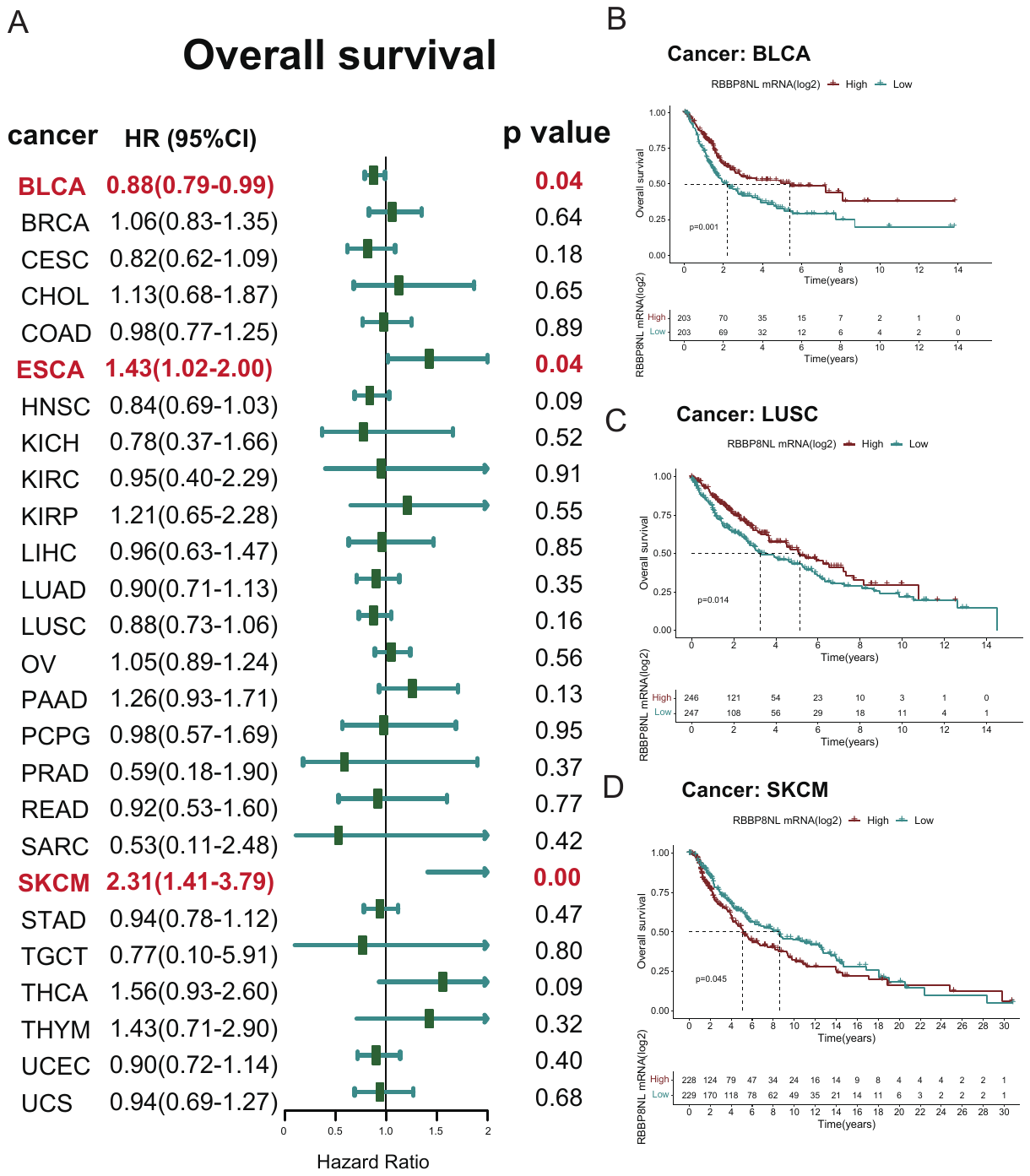

**Figure S2. Prognostic evaluation of RBBP8NL for overall survival in pan-cancers.** (A) Univariate Cox regression model used for prognostic analyses of RBBP8NL across various cancers. Hazard ratios >1 denoted risk factors, while hazard ratios <1 indicated protective factors. (B-D) Kaplan-Meier method and log-rank test were employed for prognostic analyses of RBBP8NL in pan-cancers. Only cancers where RBBP8NL emerged as a significant prognostic biomarker were included.

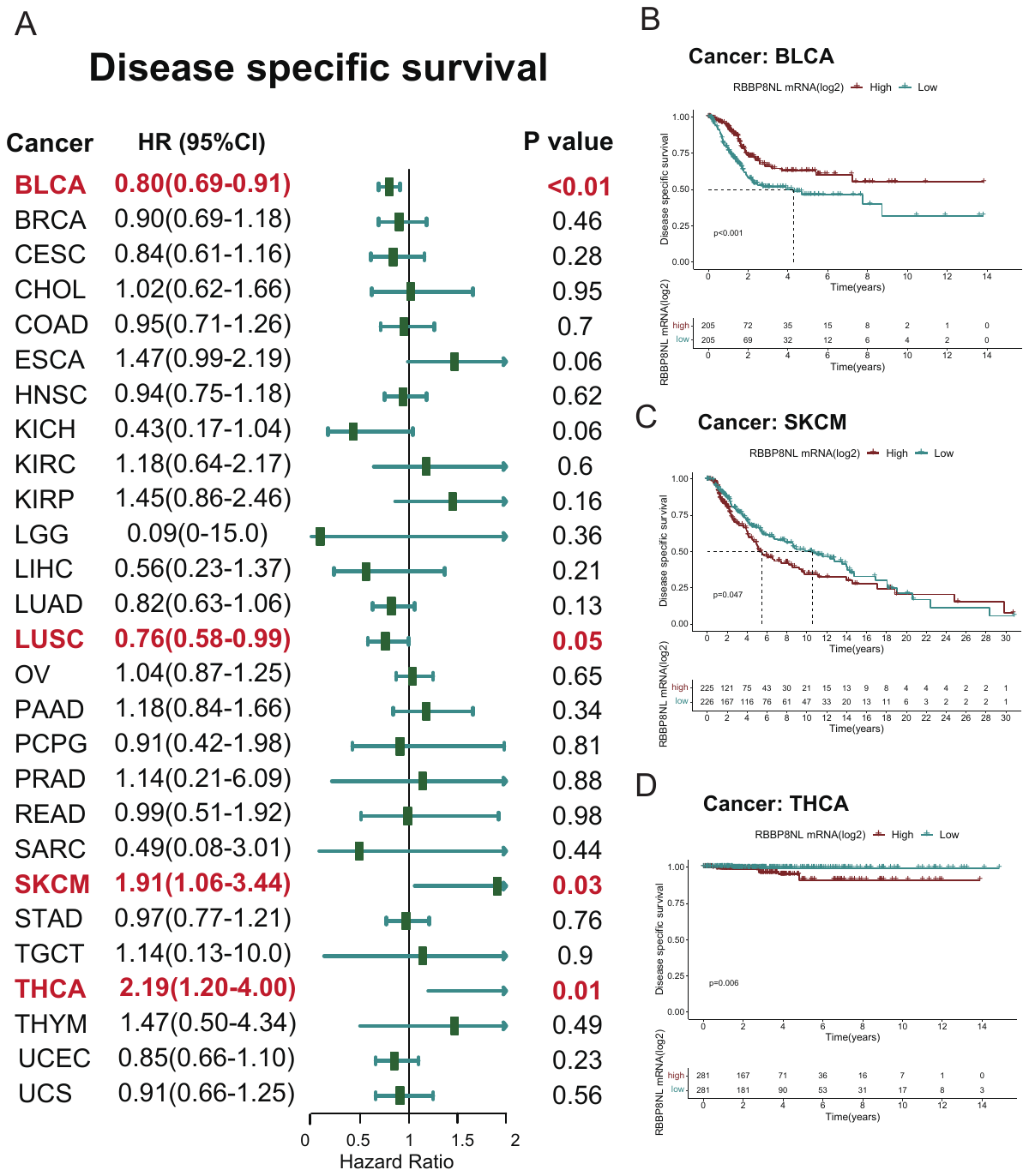

**Figure S3. Prognostic assessment of RBBP8NL for disease-specific survival in pan-cancers.** (A) Univariate Cox regression model used for prognostic analyses of RBBP8NL across various cancers. Hazard ratios >1 denoted risk factors, while hazard ratios <1 indicated protective factors. (B-D) Kaplan-Meier method and log-rank test were employed for prognostic analyses of RBBP8NL in pan-cancers. Only cancers where RBBP8NL emerged as a significant prognostic biomarker were included.

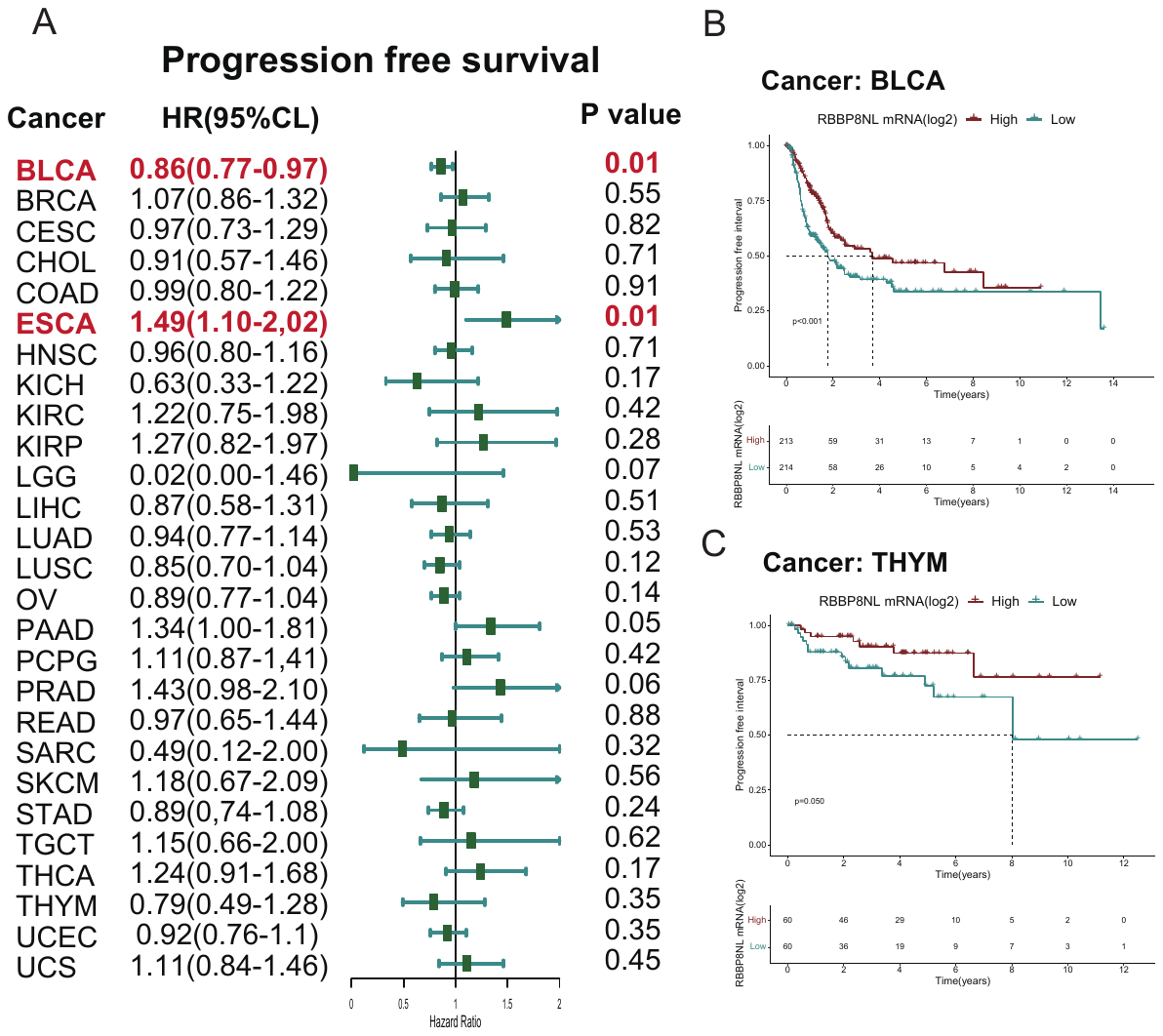

**Figure S4. Prognostic assessment of RBBP8NL for progression-free survival in pan-cancers.** (A) Univariate Cox regression model utilized for prognostic analyses of RBBP8NL across pan-cancers. Hazard ratios >1 signified risk factors, while hazard ratios <1 indicated protective factors. (B-C) Kaplan-Meier method and log-rank test were employed for prognostic analyses of RBBP8NL in pan-cancers, featuring only cancers where RBBP8NL emerged as a significant prognostic biomarker.
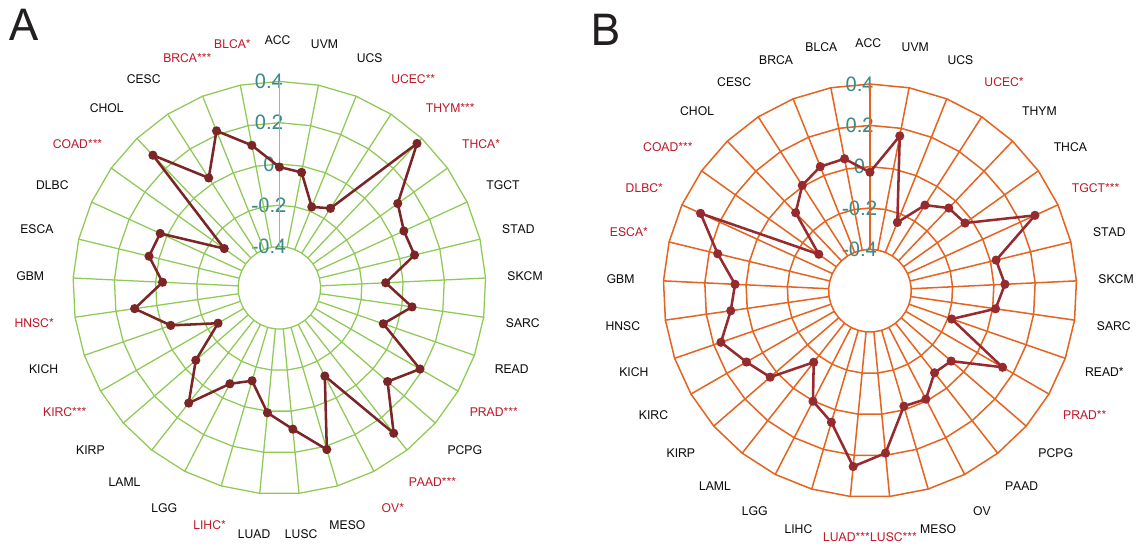

**Figure S5. Associations between RBBP8NL and TMB, MSI in pan-cancers.** (A) Relationship between RBBP8NL and TMB in pan-cancers. (B) Association between RBBP8NL and MSI in pan-cancers. The asterisks denote a significant statistical p-value determined through Spearman correlation analysis. (*P < 0.05; **P < 0.01; ***P < 0.001).
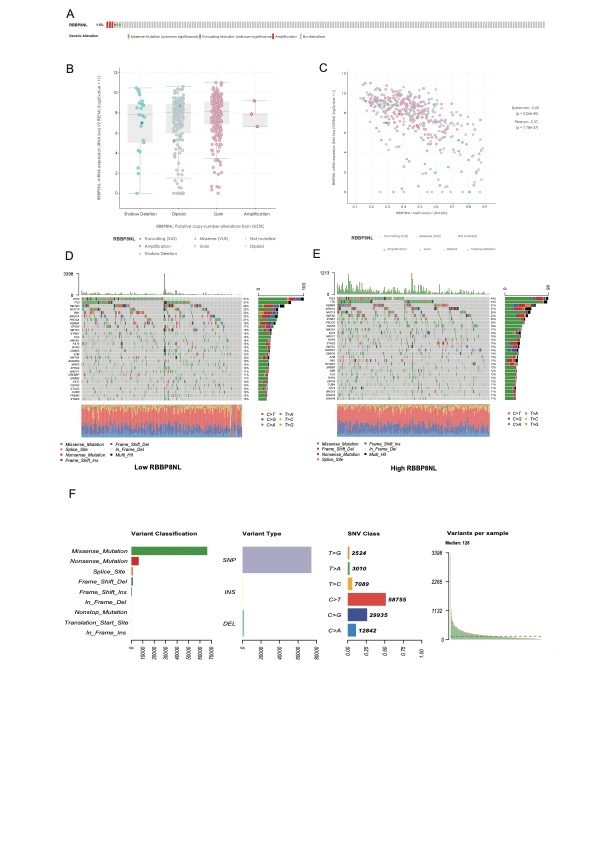

**Figure S6. Integrated analysis of RBBP8NL in BLCA.** (A) Copy number variation (CNV) landscape of RBBP8NL in BLCA. (B) Impact of RBBP8NL CNV on RBBP8NL mRNA expression. The asterisks denote a significant statistical p-value calculated with the Mann-Whitney U test (*P < 0.05; **P < 0.01; ***P < 0.001). (C) Influence of RBBP8NL methylation on RBBP8NL mRNA expression. (D-E) The top 30 mutated genes in the low and high RBBP8NL groups, respectively. The upper barplot illustrates the tumor mutation burden (TMB), with the number on the right indicating mutation frequency. The right barplot represents the distribution of variant types. (F) Overview of mutation profiles in BLCA.

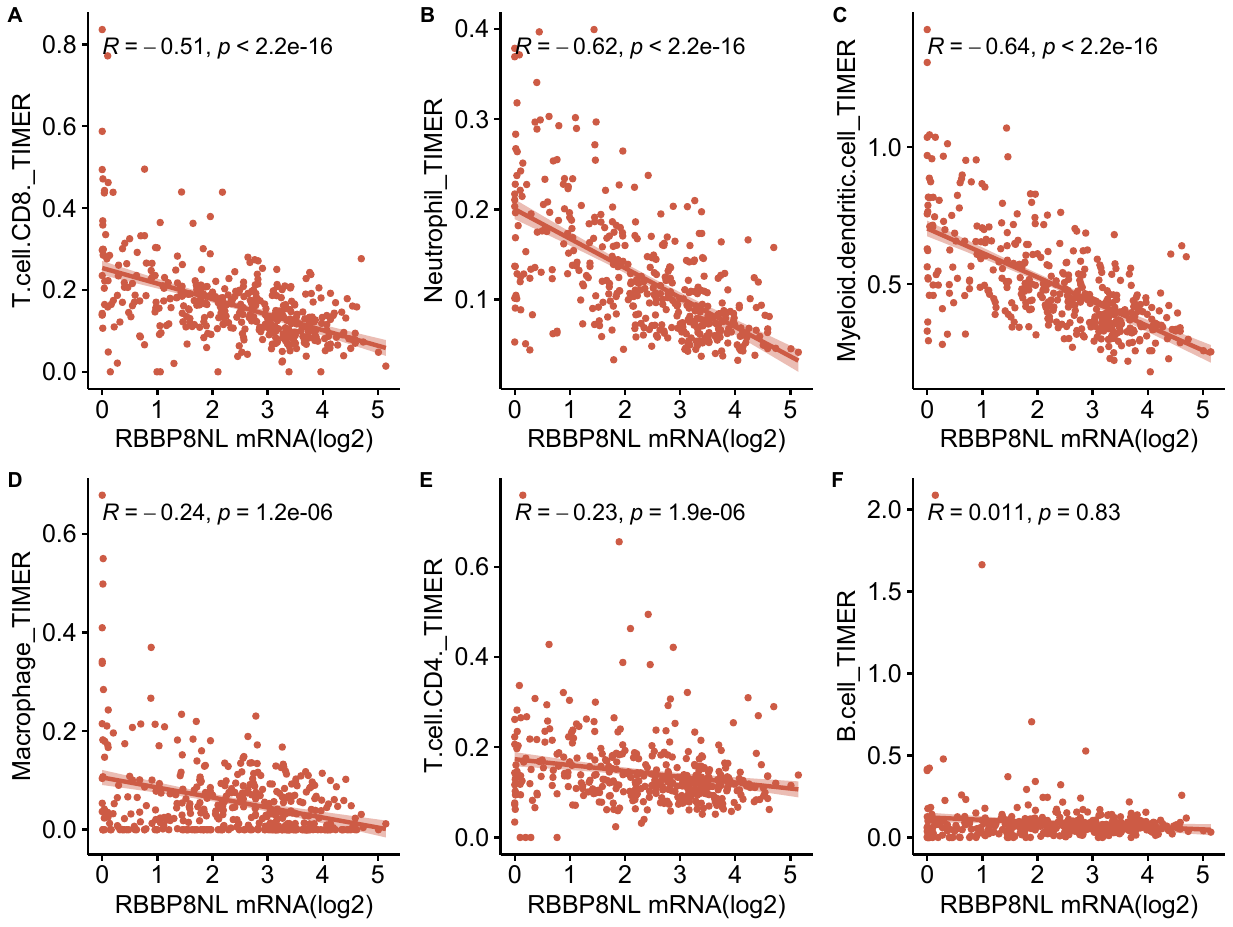

**Figure S7. Associations between RBBP8NL and immune cells in the tumor microenvironment were assessed using the TIMER algorithm.** The p-value was computed through Spearman correlation analysis.

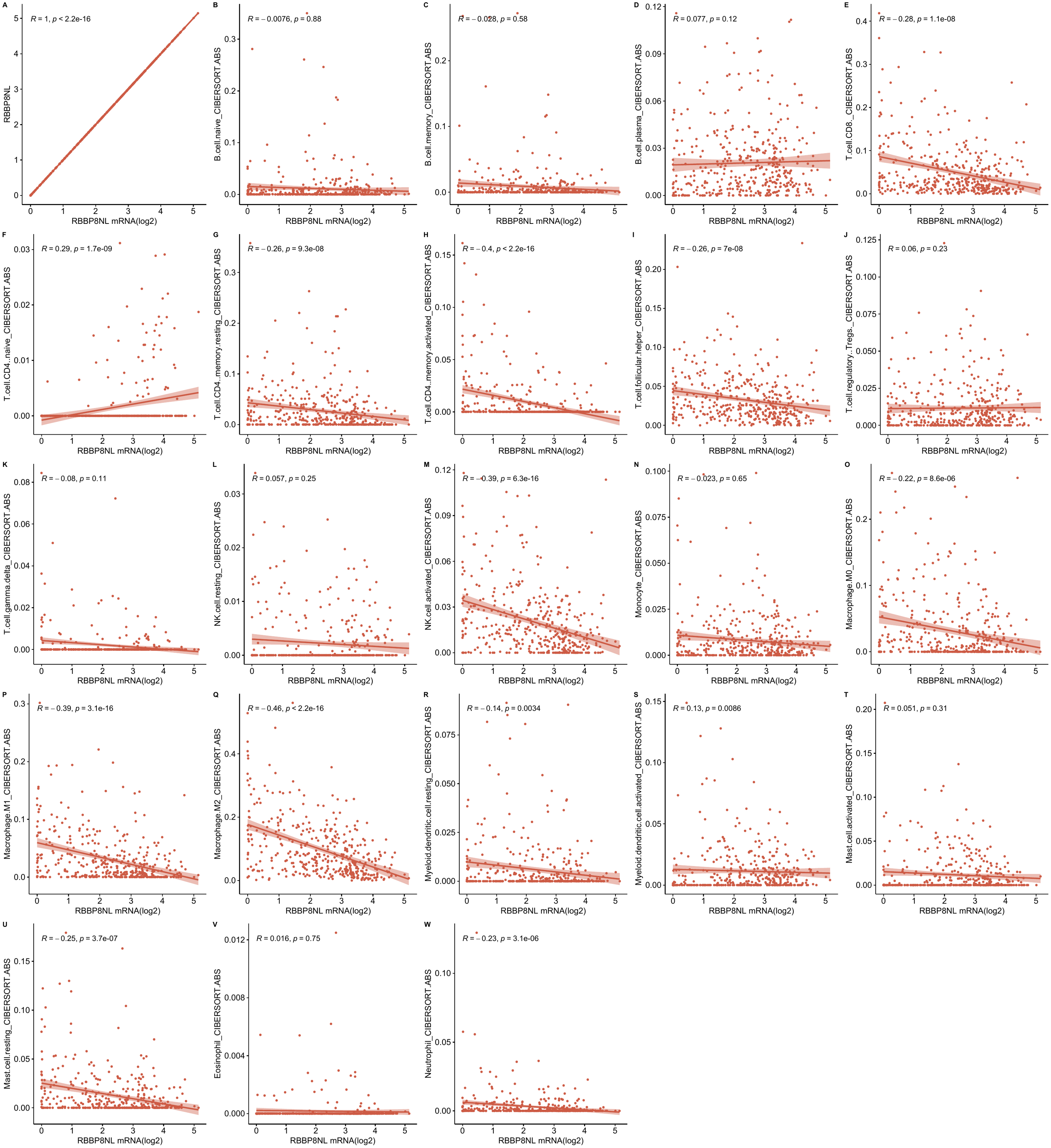

**Figure S8. Associations between RBBP8NL and immune cells within the tumor microenvironment were evaluated using the CIBERSORT-ABS algorithm.** The p-value was determined through Spearman correlation analysis.

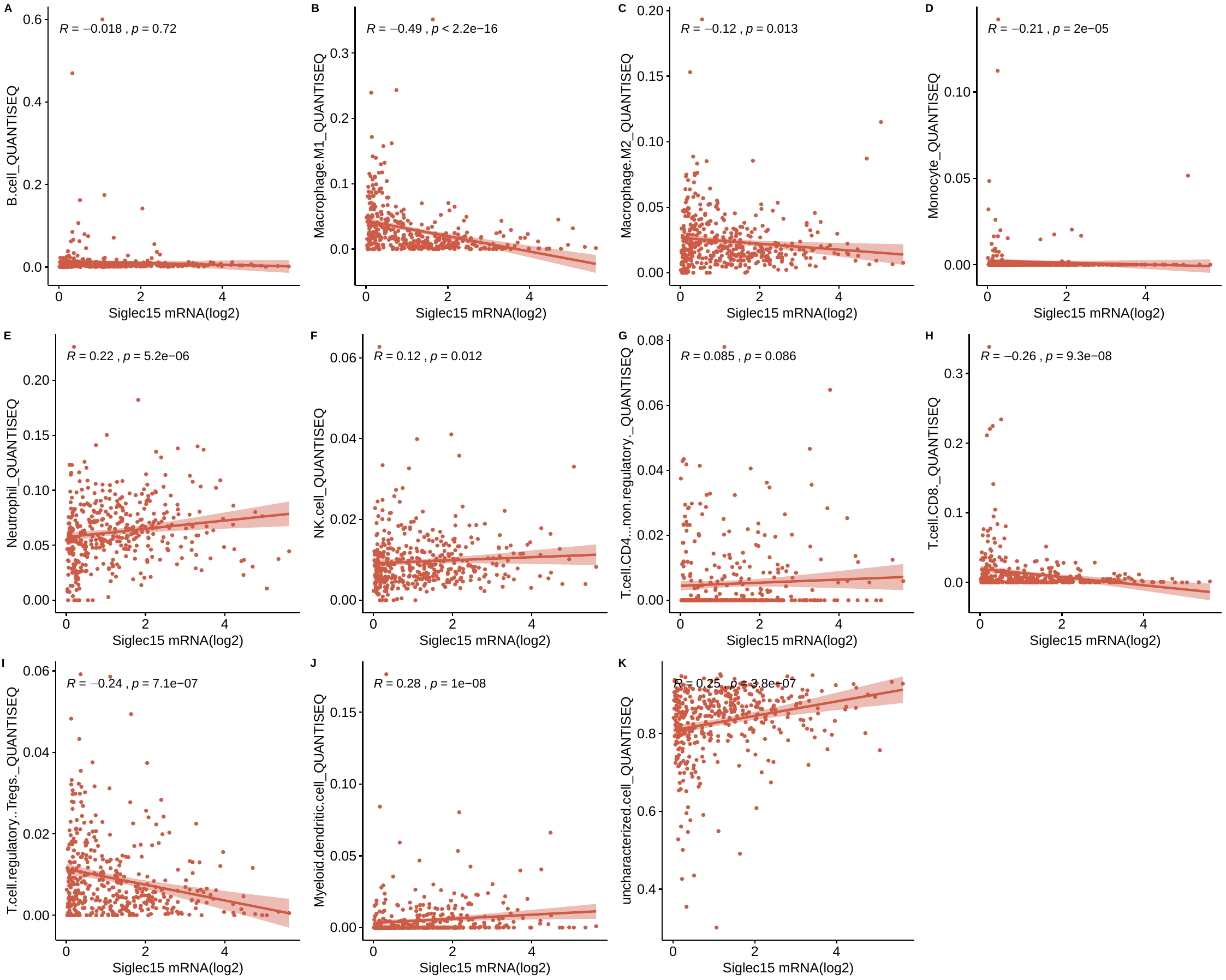

**Figure S9. Relationships between RBBP8NL and immune cells in the tumor microenvironment were assessed using the quanTIseq algorithm.** The p-value was determined through Spearman correlation analysis.

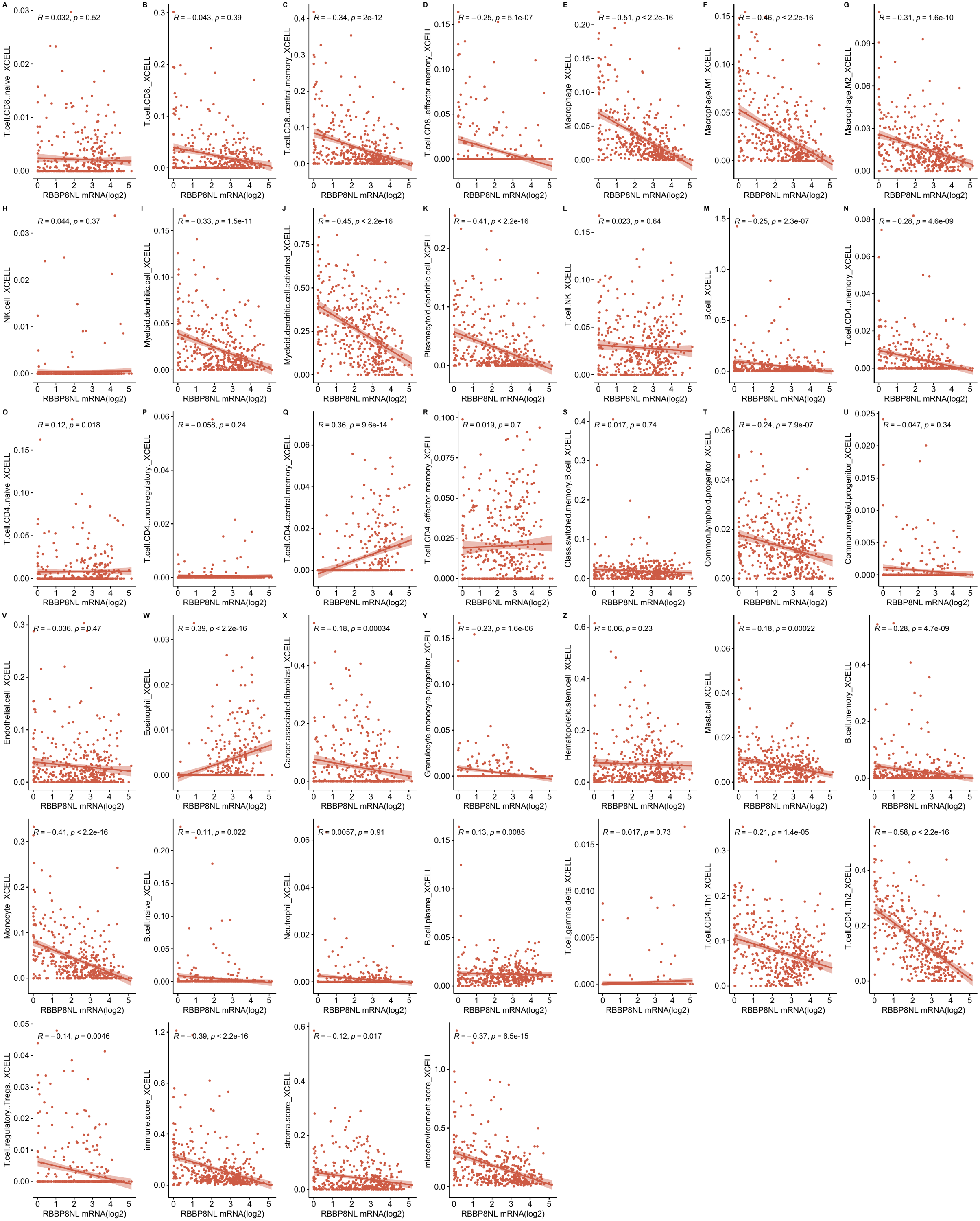

**Figure S10. Associations between RBBP8NL and immune cells within the tumor microenvironment were evaluated using the xCell algorithm.** The p-value was computed through Spearman correlation analysis.

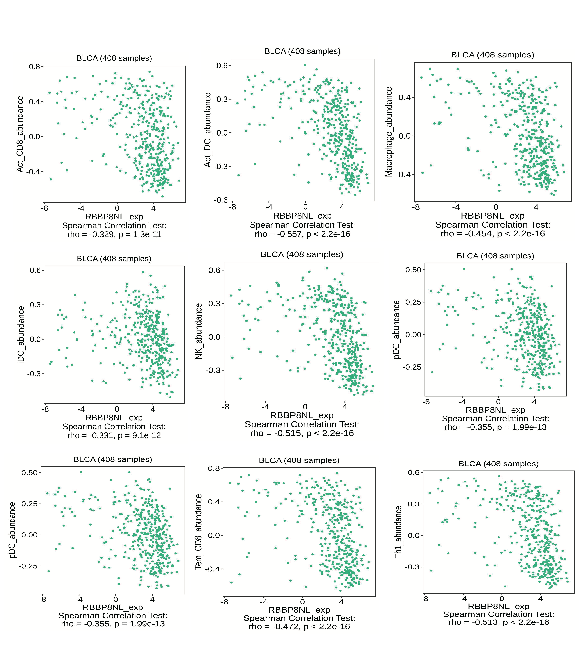

**Figure S11. Relationships between RBBP8NL and immune cells in the tumor microenvironment were assessed using the TISIDB algorithm.** The p-value was determined through Spearman correlation analysis.

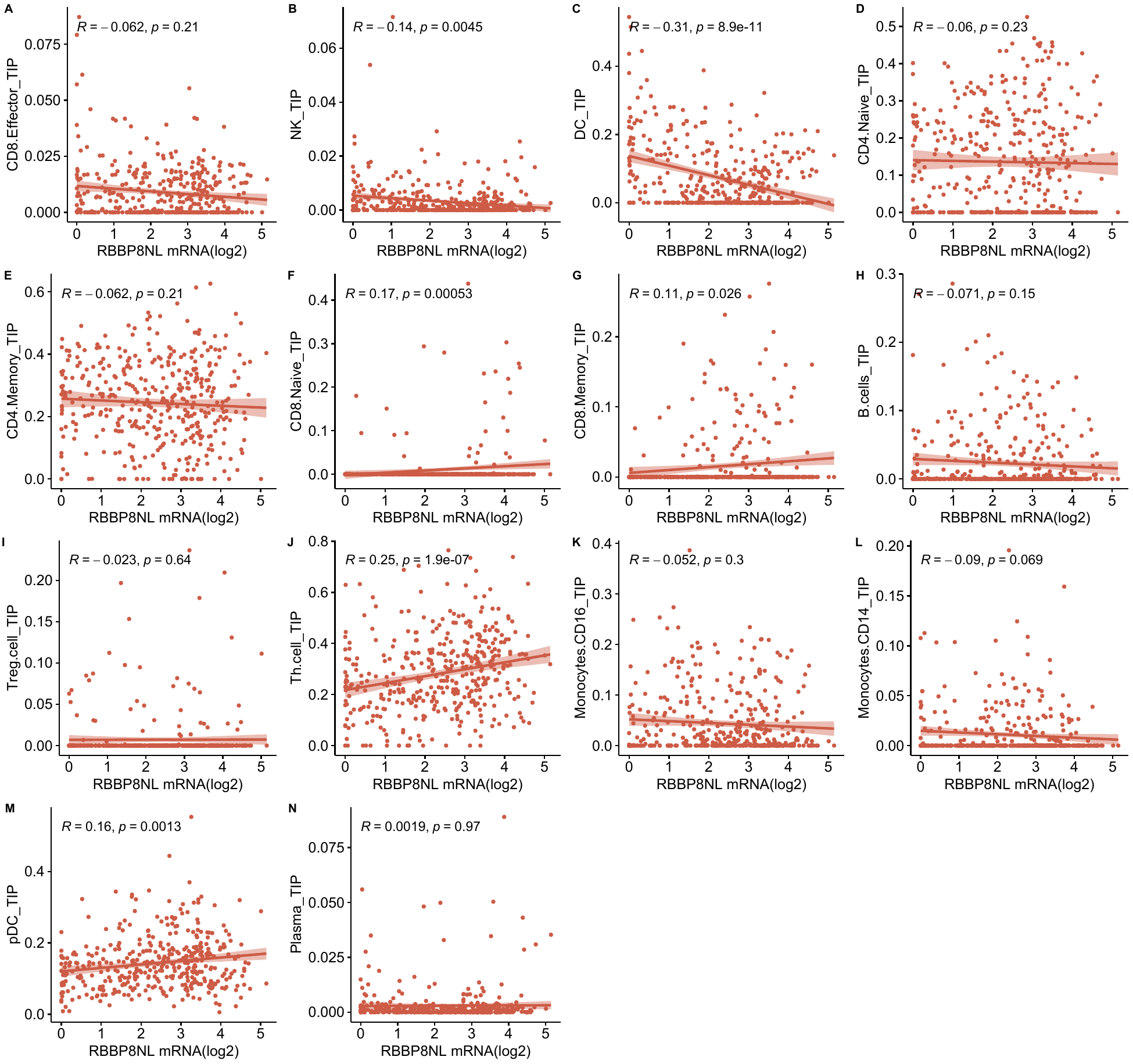

**Figure S12. Associations between RBBP8NL and immune cells in the tumor microenvironment were evaluated using the TIP algorithm.** The p-value was computed through Spearman correlation analysis.

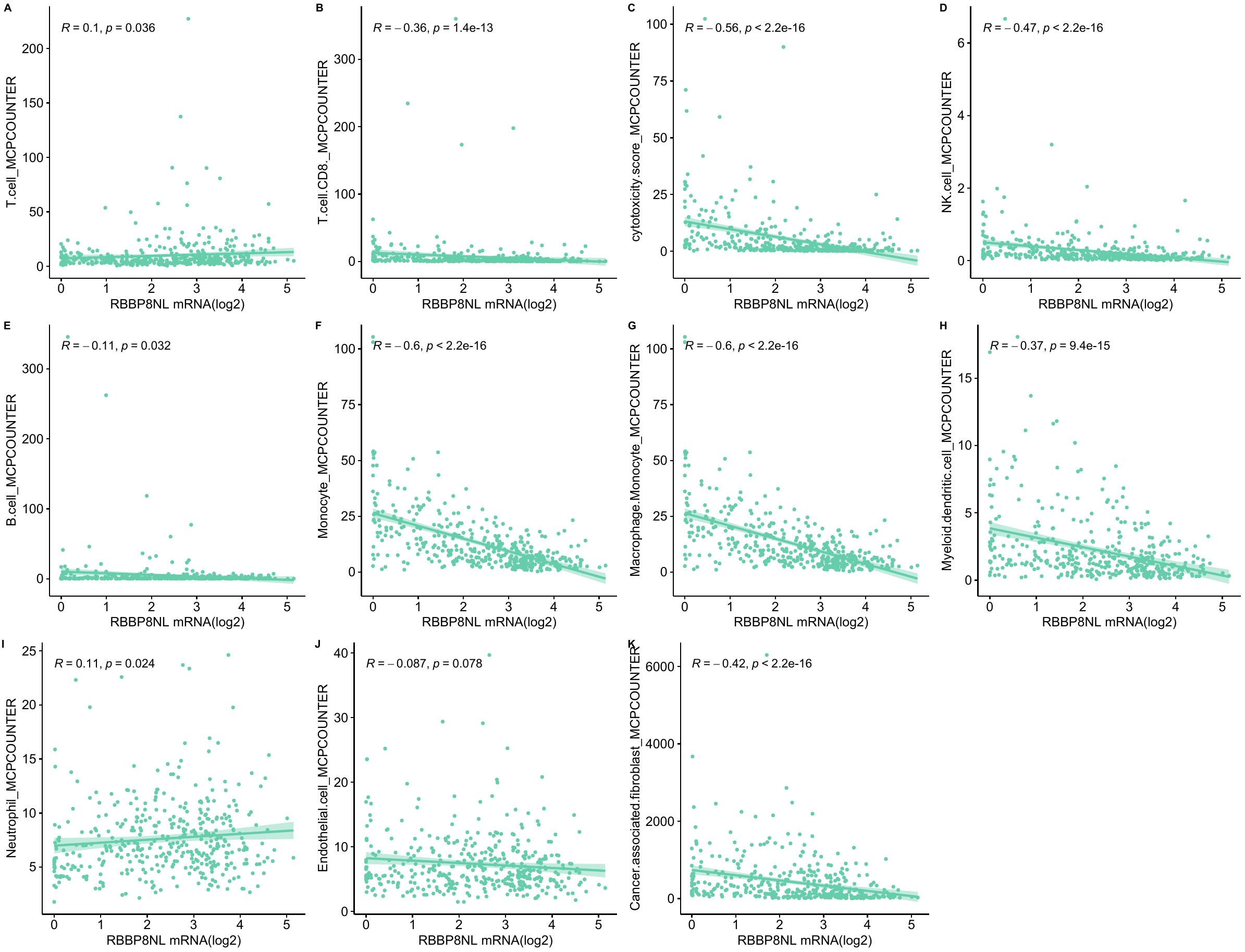

**Figure S13. Associations between RBBP8NL and immune cells in the tumor microenvironment were assessed using the MCP-counter algorithm.** The p-value was determined through Spearman correlation analysis.

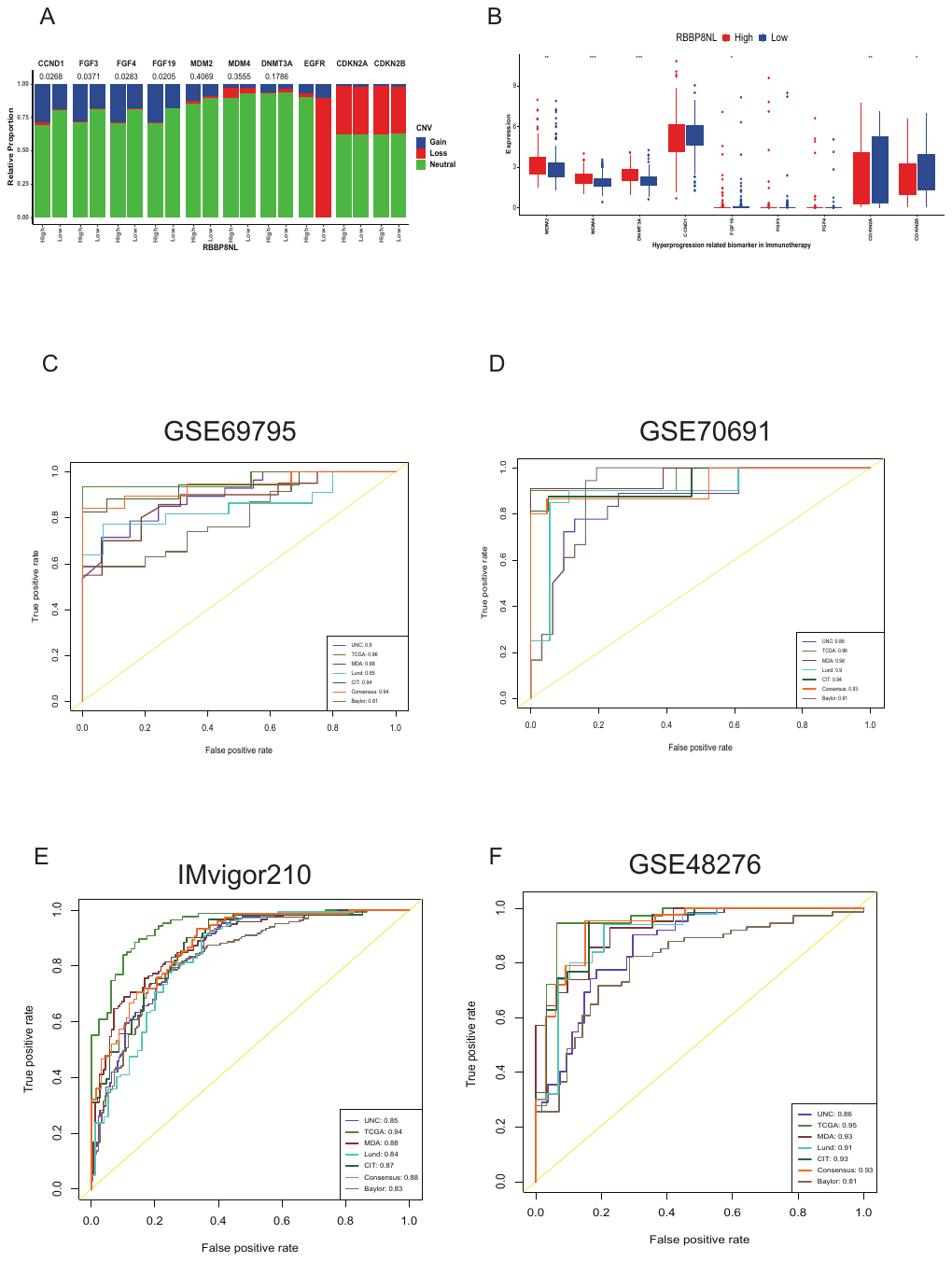

**Figure S14. Relationships between RBBP8NL and hyper-progression associated genes, and the predictive accuracy of RBBP8NL for molecular subtypes in four validation datasets.** (A) Association between RBBP8NL and CNV patterns of hyper-progression associated genes in BLCA, with the p-value determined by Fisher's t-test. (B) Correlation between RBBP8NL and mRNA expression of hyper-progression associated genes in BLCA, where significant p-values are denoted by asterisks (*P < 0.05; **P < 0.01; ***P < 0.001) using Mann-Whitney U test. (C-F) Evaluation of RBBP8NL's predictive value for molecular subtypes in four independent validation sets, comprising an immunotherapy cohort (IMvigor210), a neoadjuvant-chemotherapy cohort (GSE70691), and two general bladder cancer cohorts (GSE69795, GSE48276).
